# The autophagy activator Spermidine ameliorates Alzheimer’s disease pathology and neuroinflammation in mice

**DOI:** 10.1101/2020.12.27.424477

**Authors:** Kiara Freitag, Nele Sterczyk, Julia Schulz, Judith Houtman, Lara Fleck, Stephan J. Sigrist, Frank L. Heppner, Marina Jendrach

**Affiliations:** Department of Neuropathology, Charité – Universitätsmedizin Berlin, corporate member of Freie Universität Berlin, Humboldt-Universität zu Berlin, Berlin Institute of Health, Berlin, Germany; German Center for Neurodegenerative Diseases (DZNE) within the Helmholtz Association, Berlin, Germany; Cluster of Excellence, NeuroCure, Berlin, Germany; Freie Universität Berlin, Institute for Biology and Genetics, Berlin, Germany

**Keywords:** Alzheimer’s disease, neuroinflammation, microglia, astrocytes, autophagy, Spermidine, dietary supplement

## Abstract

Deposition of amyloid beta (Aβ) and phosphorylated Tau along with microglia- and astrocyte-mediated neuroinflammation are prominent pathogeneic features of Alzheimer’s Disease (AD). In recent years, impairment of autophagy has also been shown to contribute to AD progression. Here, we provide evidence that oral treatment of amyloid-prone AD-like APPPS1 mice with the autophagy activator Spermidine, a small body-endogenous polyamine often used as dietary supplement, decreased neuroinflammation and reduced neurotoxic soluble Aβ at both early and late stages of AD. Mechanistically, Spermidine induced autophagy in microglia as well as in astrocytes, which subsequently impacted TLR3- and TLR4-mediated inflammatory processes by decreasing cytotoxicity, inflammasome activity and NF-κB signalling. Our data highlight that autophagy targets the inflammatory response of glial cells and emphasize the potential of orally administered autophagy-activating drugs such as Spermidine to interfere with AD progression.

## Introduction

Alzheimer’s disease (AD) is the most common neurodegenerative disease and the leading cause of dementia worldwide. Due to increasing numbers of individuals affected by AD, its socioeconomic impact is enormous. In 2019, AD and dementia were among the top 10 leading causes of death according to the WHO (Global Health Estimates). Pathologically, AD is defined by three hallmarks: extracellular plaques containing amyloid-beta (Aβ) peptides, neurofibrillary tangles consisting of hyperphosphorylated microtubule-associated protein (MAP) tau, and neuroinflammation^1^. No pharmacotherapies are available to treat AD causally and as a number of recent clinical trials testing new compounds failed to improve the outcome of patients^2^, novel interventional approaches are urgently required.

Over the last decade, a large body of evidence demonstrated a substantial involvement of immune actions in AD pathogenesis. Several cytokines including interleukin (IL)-1^3^, IL-6^4^, IL-12, IL-23^5^ and tumor necrosis factor α (TNF-α)^6^ were detected in AD patients as well as in animal models with AD-like pathology^7^. Interestingly, injection of the viral mimetic PolyI:C, a synthetic analog of double-stranded RNA, into wild-type mice was sufficient for inducing an AD-related pathology^8^, pointing towards a crucial role for neuroinflammation in the onset and progression of AD. Microglia, the brain’s intrinsic myeloid cells, and astrocytes are the predominant cytokine-producing cells of the CNS. Both cell types are essential for maintaining brain homeostasis and respond to danger signals by transforming into an activated state, characterized by proliferation and cytokine release^1^. Clusters of geno- and phenotypically altered microglia and astrocytes were detected in brains of AD patients, particularly near senile Aβ plaques^9–11^. Using single cell transcriptomics, so-called disease associated microglia (DAM), which exhibit a distinct AD-associated transcriptome profile, can be clearly distinguished from homeostatic cells^12^.

A growing set of data, including those derived from GWAS studies of various human diseases by the Wellcome Trust Case Control Consortium^13^, indicates that autophagy, one of the crucial degradation and quality control pathways of the cell, can regulate inflammatory processes. Mice deficient in the autophagic protein ATG16L1 exhibit a specific increase in cytokine levels of IL-1β and IL-18 and severe colitis, which could be ameliorated by anti-IL-1β and IL-18 antibody administration^14^. Similar results were achieved in mice lacking the key autophagic factor MAP1LC3B after exposure to LPS or caecal ligation and puncture-induced sepsis^15^. Recently, we could show that a reduction of the key autophagic protein Beclin1 (BECN1), which is also decreased in AD patients^16,17^, resulted in an increased release of IL-1β and IL-18 *in vitro* and *in vivo*^18^. The mature forms of IL-1β and IL-18 are processed by activated Caspase-1 (CASP1) at the multimeric NLRP3 inflammasome complex^19^, and inflammasomes^20^ and NLRP3 aggregates^18^ were shown to be degraded by autophagy.

Autophagy efficiency seems to decline during aging^21–24^ and a growing body of evidence reveals dysfunction of autophagy in AD patients and AD model systems^25^. Autophagy presents an intriguing therapeutical target as it can be altered pharmacologically, e.g. by the small endogenous polyamine Spermidine, which is broadly used as nutritional supplement. Spermidine is known to induce autophagy by inhibiting different acetyltransferases^26,27^. Supplementation with Spermidine was found to extend the life span of flies, worms and yeast^27–30^. Moreover, Spermidine decreased the inflammatory response of macrophages and the microglial cell line BV-2 upon LPS stimulation *in vitro*^31–33^. Consistent with these observations, Spermidine supplementation improved clinical scores of and reduced neuroinflammation in mice with experimental autoimmune encephalomyelitis (EAE), a model mimicking features of multiple sclerosis^28,34^. Furthermore, Spermidine protected dopaminergic neurons in a Parkinson’s disease rat model^35^ and exhibited neuroprotective and anti-inflammatory properties in a murine model of accelerated aging^36^. These data indicate that Spermidine interferes with two dysregulated processes known to contribute to AD progression: neuroinflammation and impaired autophagy.

By administering Spermidine to AD-like APPPS1 mice we could show that soluble Aβ levels and neuroinflammation were decreased. Investigation of the involved molecular mechanism revealed that Spermidine interfered at key inflammatory signaling pathways of microglia and astrocytes, resulting in an inhibition of their activation. We therefore propose considering Spermidine as a novel therapeutic option to interfere with the progression of AD.

## Results

### Spermidine treatment of APPPS1 mice reduced soluble Aβ and neuroinflammation

Based on the previously described neuroprotective effects of Spermidine in EAE models and ageing^28,34,36–38^, we investigated the potential therapeutic effects of Spermidine on AD pathology and neuroinflammation in APPPS1 mice. This AD-like mouse model, which harbors transgenes for the human amyloid precursor protein (APP) bearing the Swedish mutation as well as presenilin 1 (PSEN1), develops a strong Aβ pathology including neuroinflammation^39^. APPPS1 mice were treated with 3 mM Spermidine via their drinking water starting prior to disease onset, namely substantial Aβ deposition, at the age of 30 days. Compared to control APPPS1 mice receiving water (H_2_O), Spermidine-supplemented animals showed no differences in fluid uptake per day (Supplementary Fig. 1a).

Mice were analyzed both at an early disease stage at 120 days and at 290 days, where pathology is known to reach a plateau^39^ (Fig. 1a). To assess Aβ deposition, consecutive protein extractions of hemispheres were performed from Spermidine-treated and control mice. Afterwards, soluble and insoluble Aβ40 and Aβ42 levels were measured by electrochemiluminescence (MesoScale Discovery panel). Strikingly, Spermidine supplementation in APPPS1 mice significantly reduced soluble Aβ40 levels in both 120 day old and 290 day old mice and the more aggregation-prone Aβ42 in 290 day old mice, while not affecting levels of insoluble Aβ (Fig. 1b-c). To substantiate these findings, we stained tissue sections for Aβ plaques using the fluorescent dye pFTAA, depicting a broad spectrum of Aβ species, and quantified the number of Aβ plaques and the average plaque size. In agreement with the MesoScale Discovery panel data, Spermidine treatment neither affected Aβ plaque covered area nor average plaque size (Fig. 1d). Also, the plaque-size distribution analysis revealed no significant differences (Supplementary Fig. 1b), indicating that Spermidine treatment preferentially targeted soluble Aβ. To gain more insights into the underlying mechanisms, we first analyzed components of the APP processing apparatus in APPPS1 mice. Western blot analysis of APP, the C terminal fragment α (CTF-α), typically generated in the non-amyloidogenic pathway, and CTF-β, a product of the amyloidogenic pathway, did not reveal any alterations, thus demonstrating that Spermidine supplementation did not influence soluble Aβ by modulating APP processing (Fig. 1e).

**Figure 1:**
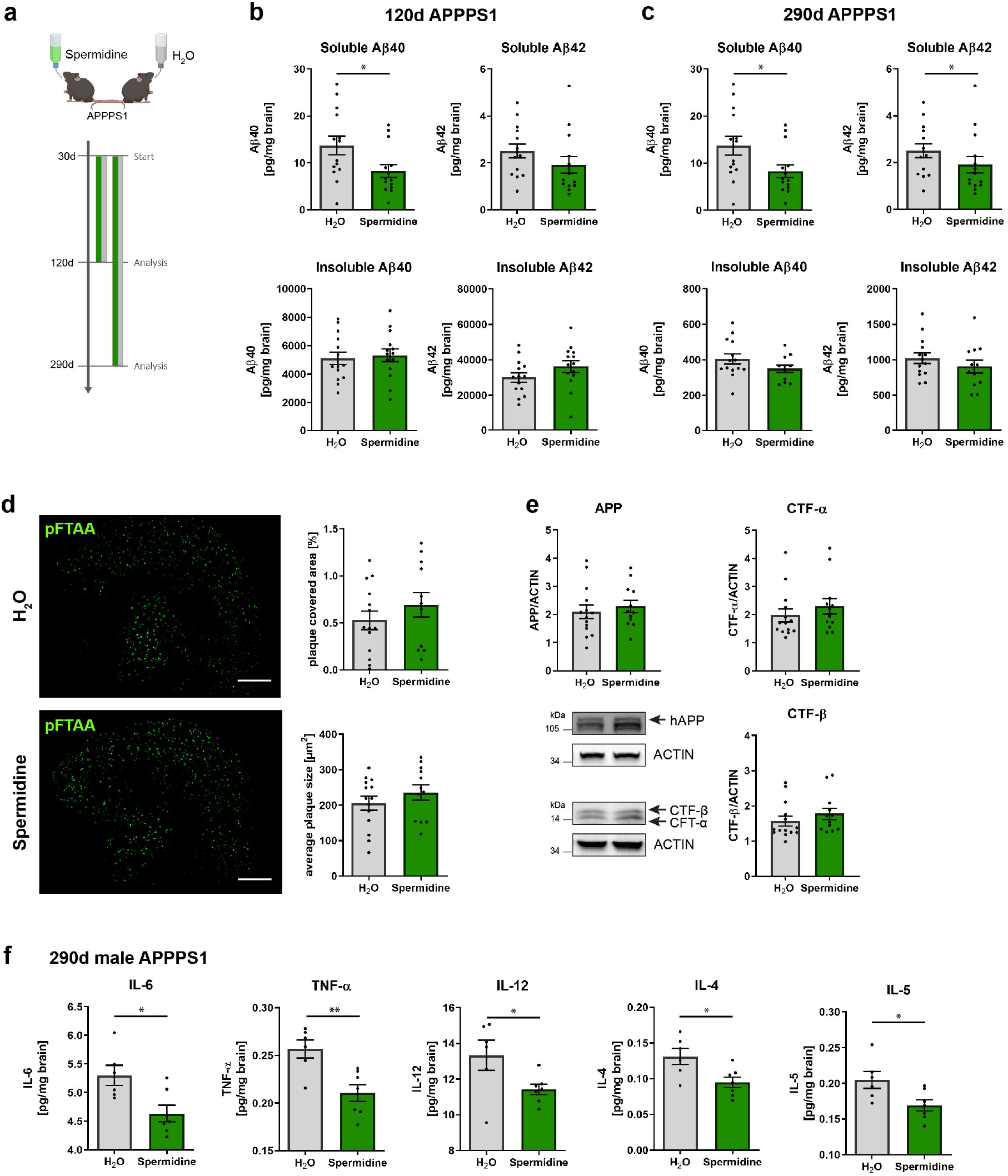
Spermidine reduced soluble Aβ and neuroinflammation in APPPS1 mice. (**a**) APPPS1 mice were treated with 3 mM Spermidine via their drinking water starting at 30 days (d) until mice reached an age of 120 days or 290 days according to the depicted treatment scheme. Spermidine-treated APPPS1 mice were compared to non-treated controls (H_2_O). (**b, c**) The Aβ40 and Aβ42 content was measured in the TBS (soluble) and SDS (insoluble) fraction of brain homogenates of (**b**) 120d old or (**c**) 290d old Spermidine-treated mice and water controls using electrochemiluminescence (MesoScale Discovery panel). 120d APPPS1 H_2_O (n = 14), 120d APPPS1 Spermidine (n = 14), 290d APPPS1 H_2_O (n = 14), 290d APPPS1 Spermidine (n = 12). (**d**) Aβ plaques in tissue sections of 290d old mice were stained with pFTAA and the plaque covered area (%) and plaque size of the cortex was determined by ImageJ analysis. APPPS1 H_2_O (n = 14), APPPS1 Spermidine (n = 12). Scale bar = 1 mm. (**e**) The levels of APP, CTF-α and CTF-β were determined by Western blot of the TX protein fraction of 290d old Spermidine-treated APPPS1 mice. Representative Western blot images are shown. APPPS1 H_2_O (n = 14), APPPS1 Spermidine (n = 12). (**f**) The content of the cytokines IL-6, TNF-α, IL-12, IL-4 and IL-5 in the TBS fraction of brain homogenates of 290d old male Spermidine-treated mice and water controls was measured using electrochemiluminescence (MesoScale Discovery panel). APPPS1 H_2_O (n = 6), APPPS1 Spermidine (n = 7). Mean ± SEM, two-tailed t-test, * p < 0.05, ** p < 0.01.

Neuroinflammation is yet another, meanwhile generally accepted, crucial regulator of Aβ pathology^1,5,40^. As neuroinflammation in AD is known to be regulated differentially depending on gender^40^, we examined the neuroinflammatory status of Spermidine-treated male APPPS1 mice. Therefore, the amount of ten cytokines were examined in brain homogenates containing the soluble proteins. Spermidine supplementation significantly reduced the AD-relevant pro-inflammatory cytokines IL-6, TNF-α, IL-12, IL-4 and IL-5 in male 290 day old mice (Fig. 1f), whilst not affecting IL-1β, IFN-γ, IL-2, IL-10 and KC/GRO (Supplementary Fig. 1c). Taken together, Spermidine treatment reduced soluble Aβ and neuroinflammation in APPPS1 mice, whilst not affecting insoluble Aβ plaques.

### Pre-treatment with Spermidine reduced TLR4-mediated inflammatory response of microglia and astrocytes

As neuroinflammatory cytokine production was affected by Spermidine treatment in APPPS1 mice *in vivo*, we focused our pathway-oriented *in vitro* analyses on microglia and astrocytes, the main cytokine-producing cells in the CNS. Therefore, microglia were isolated from adult wild type mice, pre-treated with Spermidine for 2 h and stimulated afterwards with an established protocol for activation of the TLR4 pathway (Fig. 2a). Indeed, the presence of only 30 μM Spermidine abolished the LPS/ATP-induced pro-inflammatory cytokine release of IL-1β (Fig. 2b), IL-6 and TNF-α into the cell supernatant as measured by ELISA (Supplementary Fig. 2a,b).

**Figure 2:**
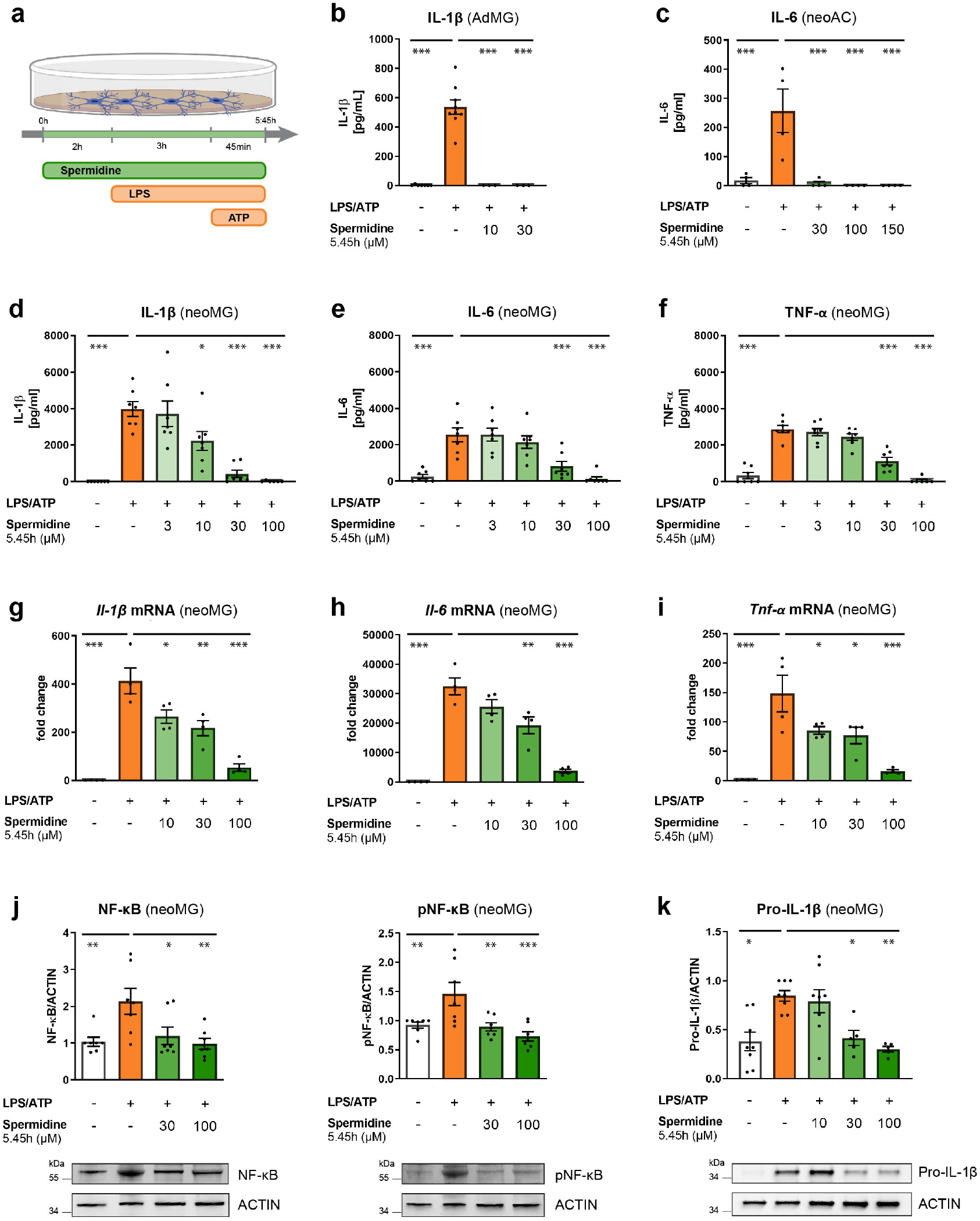
Pre-treatment with Spermidine reduced TLR4-mediated inflammatory response of microglia and astrocytes. (**a-l**) Adult microglia (AdMG), neonatal microglia (neoMG) and neonatal astrocytes (neoAC) were treated with Spermidine at indicated concentrations for 5.45 h, LPS (1 μg/ml), and ATP (2 mM) as depicted in the scheme **(a)**. (**b**) Adult microglia were isolated with magnetic activated cell sorting from 160 day old wild type mice. The concentration of IL-1β in the cell supernatant treatment was measured by ELISA. n = 3-9. (**c**) The IL-6 concentration in the cell supernatant of neonatal astrocytes (neoAC) was determined by ELISA; n = 4. (**d-f**) The IL-1β, IL-6 and TNF-α concentration in the cell supernatant of neonatal microglia (neoMG) was determined by ELISA; n = 6-7. (**g-i**) The gene expression of *Il-1β, Tnf-α* and *Il-6* was assessed by RT-qPCR. Their expression was normalized to *Actin* and displayed as fold change compared to non-treated control cells; n = 4. (**j**) Levels of phosphorylated NF-κB (pNF-κB) and NF-κB were determined by Western blot and normalized to ACTIN. Representative images are shown and protein levels are displayed as fold changes compared to non-treated controls; n = 7. (**k**) Pro-IL-1β protein levels were detected by Western blot and normalized to ACTIN. Representative images are shown and values are displayed as fold changes compared to LPS/ATP-treated cells; n = 5-8. Mean ± SEM, one-way ANOVA, Dunnett’s post hoc test (reference= LPS/ATP-treated cells), * p < 0.05, ** p < 0.01, *** p < 0.001.

To investigate the involved molecular pathways, microglia and astrocytes were isolated from neonatal wild type mice, as this model yields a higher number of cells than isolation from adult mice. Astrocytes stimulated according to the scheme depicted in Fig. 2a showed a dosedependent reduction of IL-6 in the supernatant in response to Spermidine pre-incubation (Fig. 2c), while IL-1β was released at low levels (Supplementary Fig. 2c). In neonatal microglia, treated as depicted in Fig. 2a, Spermidine treatment reduced the amount of released IL-1β, IL-6, TNF-α (Fig. 2d-f) and IL-18 (Supplementary Fig. 2d) dose-dependently, whereas IL-1β reacted most sensitively. Expanding this analysis by using electrochemiluminescence (MesoScale Discovery panel), we detected reduced levels of eight out of ten cytokines after Spermidine treatment (Supplementary Fig. 2e).

Next, we assessed whether Spermidine interfered with cytokine production or generally affected cell viability. To test the latter, lactate dehydrogenase (LDH) release as a measure of cytotoxicity was determined in neonatal microglia stimulated as depicted in Fig. 2a. While stimulation with LPS/ATP increased LDH release, addition of Spermidine dose-dependently reduced LDH (Supplementary Fig. 2f), indicating that Spermidine mediated cytoprotective effects in microglia instead of inducing cytotoxicity.

To investigate whether Spermidine can regulate the transcription of pro-inflammatory cytokines, expression of *Il-1β, Il-6* and *Tnf-α* was assessed by RT-qPCR after stimulation of neonatal microglia as depicted in Fig. 2a. In line with published data on macrophages^28^, Spermidine treatment reduced the amount of *Il-1β, Il-6* and *Tnf-α* mRNA dose-dependently (Fig. 2g-i). As Spermidine has been shown to modify the transcription factor NF-κB, responsible for the induction of *IL-1β, IL-6* and *TNF-α* expression in the cell line BV2^32^, we determined NF-κB p65 phosphorylation by Western blot. In neonatal microglia, Spermidine reduced the LPS/ATP-mediated increase in total NF-κB p65 and phosphorylated NF-κB p65 significantly (Fig. 2j).

The inflammasome, a cytosolic oligomeric signaling platform, is an a regulatory mechanism that controls IL-1β and IL-18 levels posttranslationally. Their precursors, Pro-IL-1β and Pro-IL-18, are processed by activated CASP1 at the inflammasome, which is formed after LPS/ATP stimulation and consists of NLRP3 and ASC^41^. *Nlrp3* and *Pro-Casp1* mRNA expression as well as NLRP3 protein expression were not significantly altered upon Spermidine treatment (Supplementary Fig. 2g,h). In correlation with the *Il-1β* mRNA levels (Fig. 2g), protein expression of Pro-IL-1β was reduced by administering 30 μM Spermidine (Fig. 2k). Interestingly, protein levels of Pro-CASP1 strongly increased after treatment with Spermidine, while levels of cleaved CASP1 p20 in the supernatant decreased (Supplementary Fig. 2i), hinting at a Spermidine-mediated regulation of the inflammasome activity in addition to transcriptional regulation.

### Spermidine reduced TLR3-mediated inflammatory response of microglia and astrocytes

To examine whether the anti-inflammatory effects of Spermidine were limited to TLR4-mediated inflammation, we studied its effects on the TLR3 pathway by using the pro-inflammatory stimulus PolyI:C (Fig. 3a). Confirming published observations^42,43^, PolyI:C treatment induced the release of IL-6 by adult and neonatal microglia and neonatal astrocytes, as measured in the cell supernatant by ELISA (Fig. 3b, c, d). Moreover, PolyI:C induced the release of TNF-α by adult (Supplementary Fig. 3a) and neonatal microglia (Fig. 3e), while PolyI:C-driven TNF-α release was almost absent in neonatal astrocytes (Supplementary Fig. 3d). Similar to the observed effects on the TLR4 pathway, Spermidine treatment dose-dependently reduced the PolyI:C-induced release of IL-6 and TNF-α in adult microglia, neonatal microglia and neonatal astrocytes (Fig. 3b, c, d, e, Supplementary Fig. 3a, d). While neonatal microglia already responded to low doses of Spermidine (10 μM), higher doses of 100 μM Spermidine were required to reduce the PolyI:C-driven IL-6 release in neonatal astrocytes. To exclude a putative Spermidine-induced cytotoxicity, LDH release was measured in the cell supernatant upon PolyI:C and Spermidine treatment. Based on the respective LDH levels, we concluded that Spermidine doses of up to 100 μM for neonatal microglia and 150 μM for neonatal astrocytes were non-toxic (Supplementary Fig. 3b).

**Figure 3:**
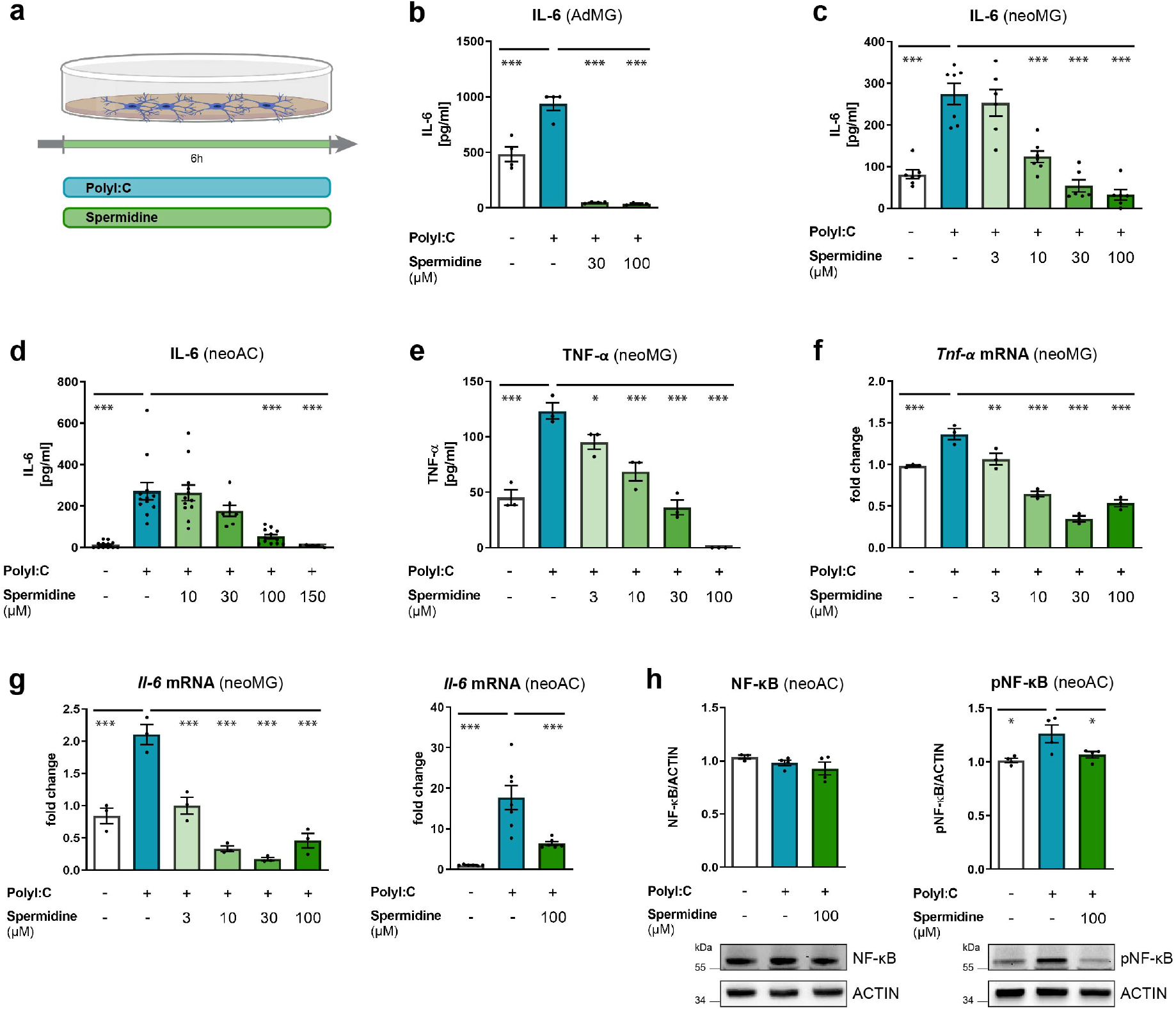
Spermidine reduced TLR3-mediated inflammatory response of microglia and astrocytes. (**a-h**) Cells were treated with PolyI:C (50 μg/ml) and Spermidine at various concentrations for 6 h as depicted in the scheme (**a**). (**b**) Adult microglia (AdMG) were isolated with magnetic activated cell sorting from 160 day old wild type mice. The concentration of IL-6 in the cell supernatant after PolyI:C and Spermidine treatment was measured by ELISA; n = 4. (**c-g**) Neonatal microglia (neoMG) and astrocytes (neoAC) were isolated from newborn mice and treated as depicted in (**a**). (**c**) The IL-6 concentration in the cell supernatant of neonatal microglia was determined by ELISA; n = 3-7. (**d**) The IL-6 concentration in the cell supernatant of neonatal astrocytes was determined by ELISA; n = 5-12. (**e**) The TNF-α concentration in the cell supernatant of neonatal microglia was determined by ELISA; n = 3-7. (**f**) The gene expression of *Tnf-α* was assessed by RT-qPCR. Its expression was normalized to *Actin* and displayed as fold change compared to non-treated control astrocytes; n = 3. (**g**) The gene expression of *Il-6* was assessed by RT-qPCR. Its expression was normalized to *Actin* and displayed as fold change compared to non-treated control microglia; n = 3-7. (**h**) Levels of phosphorylated NF-κB (pNF-κB) and NF-κB were determined by Western blot and normalized to ACTIN. Representative images are shown and values are displayed as fold changes compared to non-treated controls; n = 3-4. Mean ± SEM, one-way ANOVA, Dunnett’s post hoc test (reference= PolyI:C-treated cells), * p < 0.05, ** p < 0.01, *** p < 0.001.

To gain a broader overview of the cytokine specificity of Spermidine, we analyzed the amount of ten cytokines in the cell supernatant of neonatal microglia and astrocytes after PolyI:C and Spermidine treatment using electrochemiluminescence (MesoScale Discovery panel). Apart from IL-6 and TNF-α, also IL-12, IL-10 and KC/GRO were significantly induced by PolyI:C treatment, while they were released to a much lower extent than IL-6 (Supplementary Fig. 3c, d). The release of these cytokines was also significantly blocked by Spermidine (Supplementary Fig. 3c, d), emphazing its broad interference spectrum.

Next, we investigated whether transcriptional changes induced by Spermidine also account for the reduction in TLR3-mediated cytokine release. We found that Spermidine treatment significantly reduced PolyI:C-induced gene expression of *II-6* and *Tnf-α* in both neonatal microglia and astrocytes (Fig. 3f, g, Supplementary Fig. 3e), supporting the notion that Spermidine reduces cytokine release by interfering with the mRNA expression. To assess the involved regulatory mechanism, NF-κB p65 phosphorylation was determined by Western blot. Spermidine indeed reduced phosphorylation of NF-κB p65 in neonatal astrocytes, whilst not modulating total NF-κB levels (Fig. 3h).

Taken together, Spermidine decreased TLR4- and TLR3-mediated cytokine release in adult and neonatal microglia and neonatal astrocytes by reducing the induction of the respective mRNA expression via diminished NF-κB activity.

### Autophagy activation by Spermidine modulated the inflammatory response of glial cells

Several beneficial effects of Spermidine are attributed to Spermidine-mediated induction of autophagy^26,27^. Thus, we assessed whether Spermidine is able to induce autophagy in glial cells by probing the key autophagic factors BECN1 and LC3-II, the autophagosome-associated form of LC3 by means of Western blot. Spermidine treatment significantly upregulated BECN1 and LC3-II protein levels in LPS/ATP- and PolyI:C-treated microglia and PolyI:C-treated astrocytes (Fig. 4a,b). In agreement with a previous study, LC3-I was only weakly expressed by primary microglia^44^. Since the key transcription factor TFEB controlling autophagosomal and lysosomal biogenesis^45,46^ has been found to be regulated by Spermidine in B cells^47^, *Tfeb* mRNA expression was assessed. Indeed, while LPS/ATP and PolyI:C significantly reduced *Tfeb* mRNA in neonatal microglia and astrocytes, Spermidine treatment reversed this effect by increasing *Tfeb* to levels of non-treated cells (Fig. 4c).

**Figure 4:**
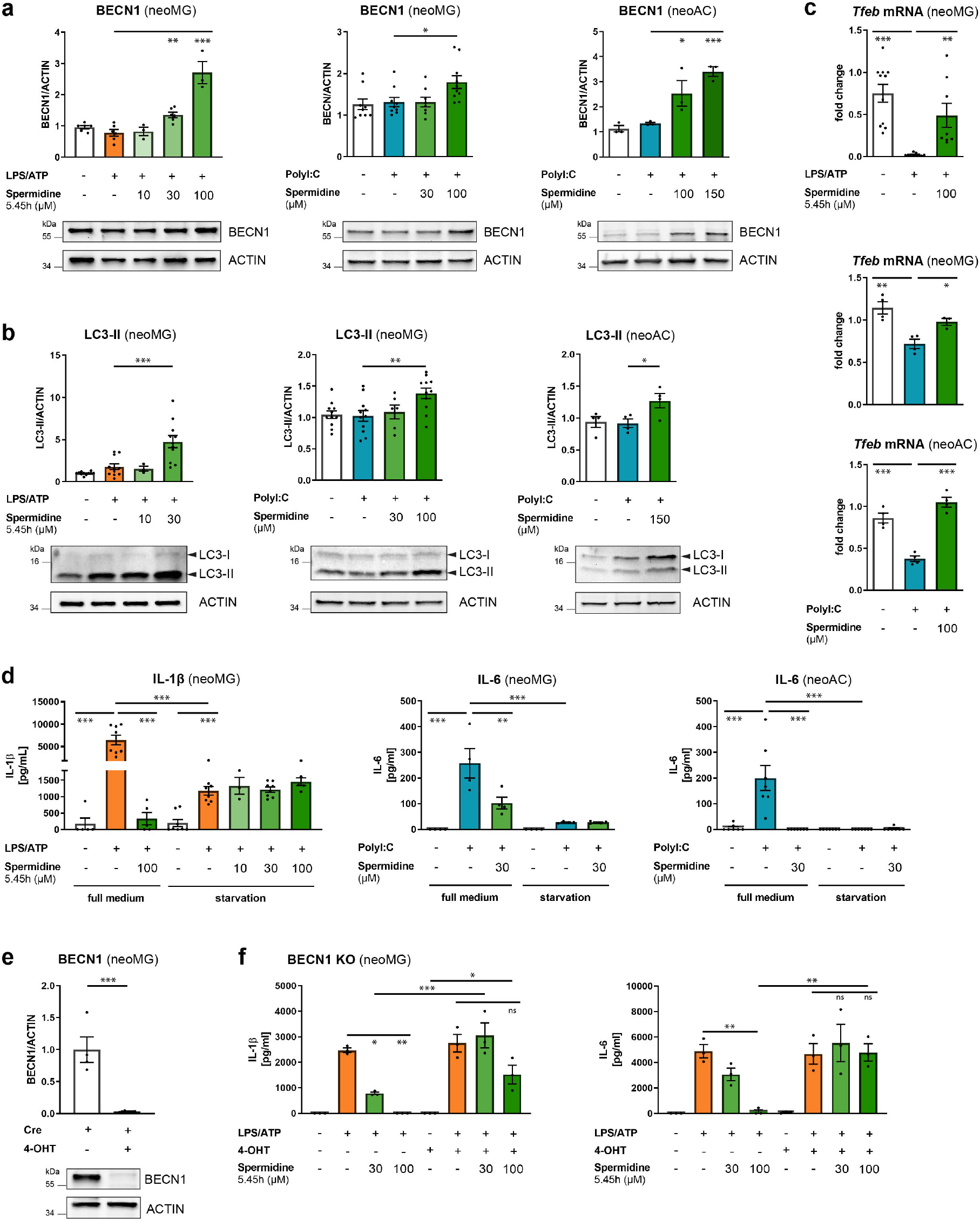
Autophagy activation by Spermidine modulated the inflammatory response of glial cells. Neonatal microglia (neoMG) and astrocytes (neoAC) were stimulated with LPS/ATP or PolyI:C and the indicated Spermidine concentrations as depicted in Fig. 2a or 3a. **(a)** BECN1 levels were determined by Western blot and normalized to ACTIN. Representative images are shown and values are displayed as fold changes compared to non-treated controls; microglia LPS/ATP n = 3-6; microglia PolyI:C n = 8-10; astrocytes PolyI:C n = 3; **(b)** The LC3-II levels were determined by Western blot and normalized to ACTIN. Representative images are shown and values are displayed as fold changes compared to nontreated controls; microglia LPS/ATP n = 3-11; microglia PolyI:C n = 6-11; astrocytes PolyI:C n = 4. **(c)** The gene expression of *Tfeb* was assessed by RT-qPCR. Its expression was normalized to *Actin* and displayed as fold change compared to non-treated control cells; microglia LPS/ATP n = 4; microglia PolyI:C n = 3-4; astrocytes PolyI:C n = 4. **(d)** Cells were stimulated either in full medium or in starvation medium HBSS and the IL-1β or IL-6 concentration in the cell supernatant was determined by ELISA; microglia LPS/ATP n = 38, microglia PolyI:C n = 4; astrocytes PolyI:C n = 7; one-way ANOVA, Tukey’s post hoc test. **(e)** Neonatal BECN1^flox/flox^.CX3CR1^Cre ERT2^ microglia were treated with Tamoxifen or Ethanol as control and BECN1 levels were assessed by Western blot and normalized to ACTIN. Representative images are shown and values are displayed as fold changes compared to nontreated controls; n = 4; two-tailed t-test. **(f)** Neonatal BECN1^flox/flox^.CX3CR1^Cre ERT2^ microglia were treated with Tamoxifen (4-OHT) or Ethanol as control and subsequently treated as depicted in Fig. 2a. The release of IL-1β and IL-6 was determined by ELISA; n = 3. Mean ± SEM, one-way ANOVA, Dunnett’s post hoc test (reference = LPS/ATP or PolyI:C-treated cells) if not stated otherwise, * p < 0.05, ** p < 0.01, *** p < 0.001.

To determine whether autophagy induction is crucial for the anti-inflammatory action of Spermidine, cells were treated with LPS/ATP and Spermidine as shown in Fig. 2a, and with PolyI:C and Spermidine as depicted in Fig. 3a. Here, either full medium or amino acid-free starvation medium HBSS, known to achieve a robust induction of autophagy, was used. As shown previously^18^, cell starvation impaired the release of cytokines upon LPS/ATP or PolyI:C treatment significantly (Fig. 4d), underlining that autophagy induction reduces the inflammatory response of glial cells. In addition, no further reduction of cytokine release after Spermidine treatment became discernable.

In order to assess whether reduction of autophagy can prevent Spermidine-mediated cytokine decrease, we performed the reverse experiment by genetically deleting BECN1 using a Cre-flox approach. To induce excision of the floxed *Becnl* gene gene, we treated neonatal microglia derived from BECN1^flox/flox^.CX3CR1^CreERT2^ mice with Tamoxifen or Ethanol as a control, ultimately resulting in a strong reduction of BECN1 protein expression in Tamoxifen-treated cells (Fig. 4e). Subsequently, cells were stimulated with LPS/ATP and Spermidine as depicted in Fig. 2a. Quantification of the IL-1β and IL-6 release by ELISA showed a significantly decreased sensitivity of BECN1-deficient microglia to Spermidine (Fig. 4f). Taken together, these data confirm that Spermidine induces autophagy in glial cells, thus mediating the reduction of cytokine release.

### Spermidine modulated the IL-1β processing pathway in activated microglia

After establishing that pre-treatment and simultaneous treatment with Spermidine ameliorated the pro-inflammatory response of glial cells, we subsequently investigated whether Spermidine could excert its anti-inflammatory action in previously activated microglia, thus contributing to the reduction of soluble Aβ and cytokines in the Spermidine-treated APPPS1 mice (Fig. 1b, c, f). Therefore, microglia were first primed for 2 h with LPS before adding Spermidine (Fig. 5a). Indeed, even post-activation treatment of primed adult (Fig. 5b) and neonatal microglia (Fig. 5c) with Spermidine reduced IL-1β and IL-18 (Supplementary Fig. 4a) release into the cell supernatant dose-dependently. Interestingly, IL-6 and TNF-α were not altered by this Spermidine administration protocol (Supplementary Fig. 4b). We expanded our analysis by using electrochemiluminescence assays and found that in fact only IL-1β and IL-18 were regulated by Spermidine but none of the other nine measured cytokines (Supplementary Fig. 4c).

**Figure 5:**
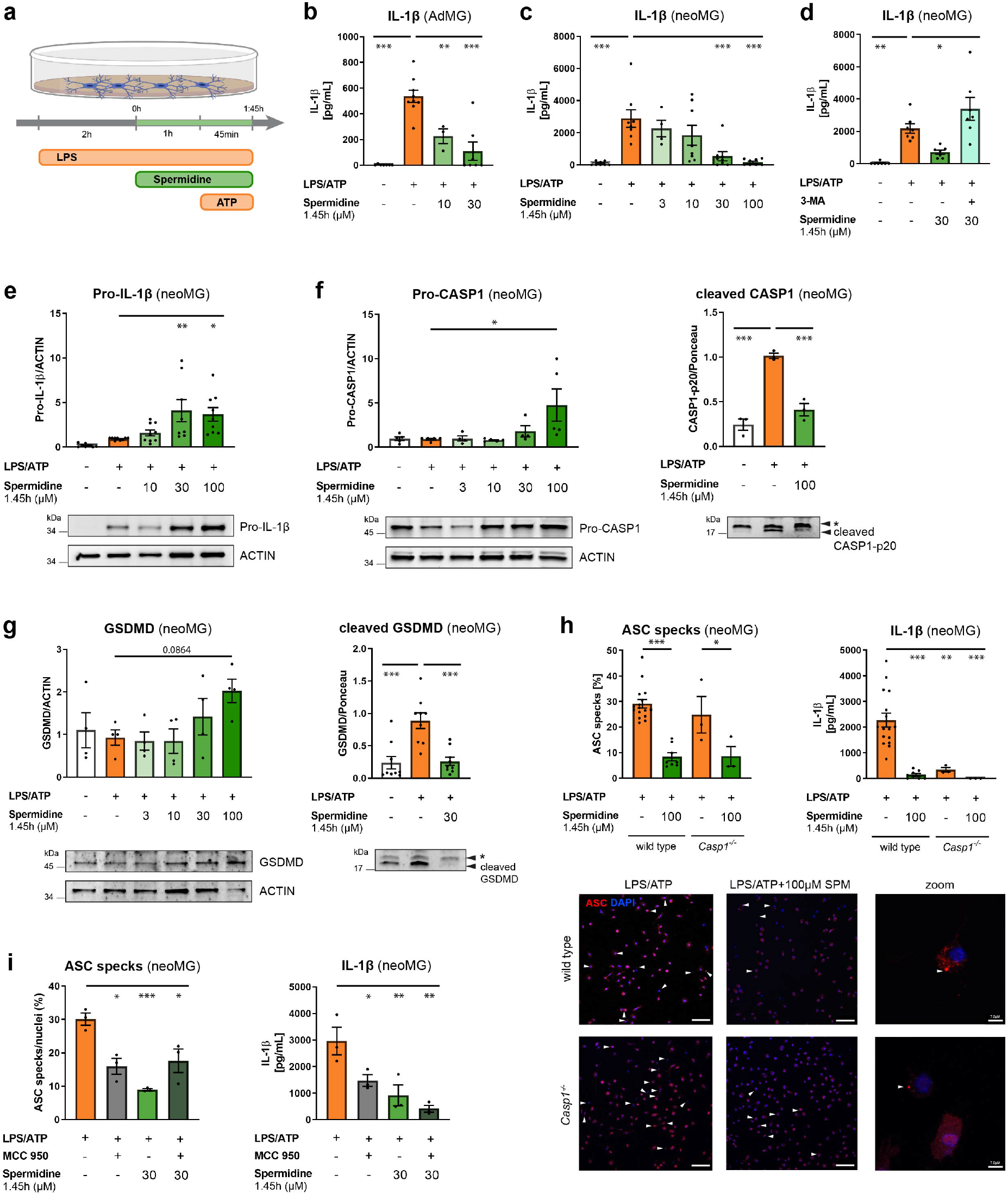
Spermidine modulated the IL-1β processing pathway in activated microglia. (**a**) Cells were treated with LPS (1 μg/ml) and Spermidine at indicated concentrations for 1.45 h and ATP (2 mM) as depicted in the scheme. **(b)** Adult microglia (AdMG) were isolated with magnetic activated cell sorting from 160 day old wild type mice. The concentration of IL-1β in the cell supernatant after LPS/ATP and Spermidine treatment was measured by ELISA. Non-treated and LPS/ATP-treated data points are the same as in Fig. 2b; n = 3-9. **(c-i)** Neonatal microglia (neoMG) were treated with LPS/ATP and Spermidine as shown in (**a**). **(c)** The IL-1β concentration in the cell supernatant was determined by ELISA; n = 4-8. **(d)** Neonatal microglia (neoMG) were stimulated as shown in (**a**) and 3-MA was added simultaneously with Spermidine. The IL-1β concentration in the cell supernatant was determined by ELISA; n = 7. **(e)** Pro-IL-1β protein levels were determined by Western blot and normalized to ACTIN. Representative images are shown and values are displayed as fold changes compared to LPS/ATP-treated cells; n = 8-9. **(f)** Cellular Pro-CASP1 levels protein levels and cleaved CASP1 levels in the supernatant were determined by Western blot (* non-specific band). Pro-CASP1 was normalized to ACTIN (n = 4-8) and CASP1 was normalized on whole protein content determined by Ponceau S staining (n = 3). Values are displayed as fold changes compared to LPS/ATP-treated cells. **(g)** Cellular and cleaved GSDMD (C-terminal fragment) levels in the supernatant were determined by Western blot (* non-specific band). GSDMD was normalized to ACTIN (n = 4) and the C-terminal fragment was normalized to whole protein content determined by Ponceau S staining (n = 9). Values are displayed as fold changes compared to LPS/ATP-treated cells. **(h)** Neonatal wild type and *Casp1^-/-^* microglia were stimulated as shown in (**a**) but with 4 mM ATP to increase the number of inflammasomes. Cells were stained for ASC (red) to visualize inflammasomes and with DAPI (blue) for nuclear staining as shown in the representative images (scale bar = 100 μm). The percentage of ASC specks in respect to the number of total cells (DAPI positive cells) was determined (left). One-way ANOVA, Tukey’s post hoc test. The IL-1β concentration in the cell supernatant was assessed by ELISA (right); wild type: n = 8-16; *Casp1^-/-^*: n = 3. **(i)** Neonatal microglia were stimulated as shown in (**a**) and MCC950 was added 15 min before addition of ATP. Cells were stained for ASC to visualize inflammasomes and with DAPI for nuclear staining. The percentage of ASC specks in respect to the number of total cells (DAPI positive cells) was determined (left). The IL-1β concentration in the cell supernatant was assessed by ELISA (right); n = 3. Mean ± SEM, one-way ANOVA, Dunnett’s post hoc test (reference = LPS/ATP-treated cells) if not stated otherwise, * p < 0.05, ** p < 0.01, *** p < 0.001.

To assess the status of autophagy, Western blot analyses of LC3-II showed that even a short Spermidine treatment of 1.45 h was sufficient to activate autophagy (Supplementary Fig. 4d). To learn whether autophagy activation also mediated the reduction of IL-1β in this setting, cells were treated according to Fig. 5a in the amino acid-free starvation medium HBSS. Induction of autophagy by HBSS reduced IL-1β release, as shown in Fig. 4d and again, no additional decrease of cytokine release in Spermidine-treated cells was observed (Supplementary Fig. 4e). For the complementary approach, cells were treated according to the scheme in Fig. 5a and the autophagy-inhibitor 3-MA was added simultaneously with Spermidine. Addition of 3-MA abolished the Spermidine-mediated reduction of IL-1β release completely (Fig. 5d). In contrast, IL-6 and TNF-α expression were not changed by 3-MA treatment (Supplementary Fig. 4f), supporting the conclusion that Spermidine reduces IL-1β release in activated microglia in an autophagy-dependent manner.

Measurement of LDH release showed that Spermidine was cytoprotective in this setting as well (Supplementary Fig. 4g), implying that Spermidine may also exert an anti-inflammatory action by reducing pyroptosis and thus impairing the release of IL-1β and IL-18 from the cell.

While Pro-IL-1β protein levels beginning at 30 μM Spermidine were significantly increased (Fig. 5e), *Il-1β* mRNA levels were not altered (Supplementary Fig. 4h). These data indicate that Spermidine also interfered with the IL-1β pathway on the protein level. Thus, we investigated the key steps of the IL-1β processing pathway. Starting with the precursor of active CASP1, increased levels of uncleaved Pro-CASP1 in microglia treated with 100 μM Spermidine were detected, correlating with the observed reduction of cleaved/activated CASP1 in the supernatant (Fig. 5f). This effect was congruent with the results obtained after 5.45 h Spermidine treatment (Supplementary Fig. 2i). In agreement with this observation, cleavage of Gasdermin D (GSDMD), another substrate of CASP1, was also strongly reduced in Spermidine-treated microglia (Fig. 5g). Upon observing that the precursors of the IL-1β processing pathway accumulated while the active forms were reduced, we investigated the effects of Spermidine on the inflammasome. NLRP3 levels were not altered on the mRNA or protein level by Spermidine (Supplementary Fig. 4i). However, staining and quantification of ASC specks/inflammasomes, the processing sites of Pro-CASP1, GSDMD and Pro-IL-1β, revealed that adding Spermidine reduced the number of ASC specks significantly (Fig. 5h). A similar reduction was also detected in *Casp1^-/-^* microglia, indicating that Spermidine did not directly interfere with Pro-CASP1 cleavage but rather with the inflammasome formation. To test this hypothesis, the ASC-oligomerization inhibitor MCC950^48^ was added 15 min before addition of ATP (Fig. 5a). Since we detected no additive effects of MCC950 in regard to IL-1β release or number of ASC specks beyond that of Spermidine, one can assume that Spermidine is indeed interfering with ASC-oligomerization and with the formation of inflammasomes in activated microglia (Fig. 5i). Consistent with this hypothesis, Western blot analyses for ASC after chemical crosslinking showed reduced appearance of ASC oligomers in Spermidine-treated cells (Supplementary Fig. 4j), while the amount of ASC monomers was not altered (Supplementary Fig. 4k). Taken together, these data reveal that Spermidine treatment reduced the processing of IL-1β in activated microglia by interfering with the oligomerization of ASC-positive inflammasomes.

All in all, our data substantiate the potential of Spermidine to modulate the imperative processes and mechanisms of glial inflammation during AD. Spermidine reduced TLR3- and TLR4-mediated activation of microglia and astrocytes via distinct molecular mechanisms, thus targeting essential Aβ-regulating pathways.

## Discussion

Targeting AD pathology by preventing or at least delaying disease progression has been and remains challenging, representing an urgent unmet clinical need. Based on recent advances in our understanding of AD pathogenesis resulting in the appreciation of the impact of neuroinflammation and autophagy processes, we assessed the therapeutic effects of the autophagic activator Spermidine in the AD-like mouse model APPPS1. Strikingly, Spermidine treatment significantly reduced soluble Aβ levels at 120 days (early pathology) and at 290 days (late pathology), while Aβ plaque burden and size were not altered. Currently, it is thought that soluble Aβ causes more synaptotoxicity than plaque-bound insoluble Aβ by altering synaptic transmission and mediating synaptic loss and neuronal death, thus stressing the importance of targeting soluble Aβ in AD^49–51^. As the APPPS1 mouse model exhibits a fast disease progression with a strong genetically-driven amyloid pathology starting already at 60 days^39^, the effect of Spermidine on soluble Aβ hints at the potential of Spermidine to counteract or at least slow AD progression.

Mechanistically, we can show that Spermidine did not alter APP processing *in vivo*, supporting the notion that the regulation of glial activation and neuroinflammation appears to be a major mechanism in driving the reduction of soluble Aβ. Indeed, we observed a substantial reduction of neuroinflammation in 290 day old APPPS1 mice. Consistent with our observations, elevated pro-inflammatory cytokine production has been shown to trigger accumulation of Aβ species *in vivo*, and Aβ production in neurons *in vitro*^4,8,52–54^. Furthermore, pro-inflammatory cytokine production was decreased and disease pathology was ameliorated in Spermidine-treated EAE mice, a murine model for multiple sclerosis^28,34^. However, in the latter study, Spermidine was injected intraperitoneally, while we administered Spermidine orally, which is yet another substantial benefit of our strategy in light of the possible applicability of Spermidine in humans.

To unravel the mechanism of Spermidine-mediated effects on neuroinflammation in APPPS1 mice, we deduced the complexity of the *in vivo* setting to the main cytokine-producing glial cells, namely microglia and astrocytes. By modelling neuroinflammation *in vitro* through activation of the TLR3 and TLR4 pathway, we found that Spermidine treatment abolished the release of pro-inflammatory cytokines by primary adult and neonatal microglia and neonatal astrocytes. These results indicate that Spermidine targets both glial cell types, likely explaining the observed reduction of neuroinflammation *in vivo*. In support of our data, Spermidine treatment has been reported to reduce LPS-induced release of IL-6, TNF-α, IL-1β and IL-12 in dendritic cells, macrophages and the microglial cell line BV2^32,55–57^. Moreover, beyond what we observed in Spermidine-treated microglia, the neuroinflammatory response of astrocytes was altered upon exposure to Spermidine, thus illustrating a broad spectrum of cells affected by Spermidine.

With respect to the mode of Spermidine action, NF-κB has been shown to be involved in regulating transcription of pro-inflammatory genes in Spermidine-treated BV2 and dendritic cells^32,57^. In line with these reports we also found that Spermidine regulated the response of primary glial cells through NF-κB. By extending these previous findings we can additionally show that Spermidine actions were not limited to the TLR4 pathway but also inhibited TLR3-mediated inflammation. Of note, Spermidine reduced the LPS/ATP-induced release of IL-1β with high specificity also in a posttranslational manner by interfering with the assembly of ASC-containing inflammasomes in primed microglia, thus indicating that Spermidine action goes beyond regulation of transcripts. We observed no changes in IL-1β levels *in vivo*, which may be due to the fact that the MesoScale Discovery panel - which we used to assess IL-1β - does not discriminate between Pro-IL-1β and mature IL-1β (personal communication with MesoScale Discovery). Noteworthy, inhibiting the IL-1β processing NLRP3 inflammasomes, either by genetic knockout or pharmacologically by the inhibitor MCC950, was described to improve Aβ clearance and spatial memory in an Aβ-harbouring mouse model, and was shown to reduce α-synuclein pathology and neurodegeneration in Parkinson’s disease mice, respectively^58,59^.

Another mode of action of Spermidine was its cytoprotective effect as shown by reduced release of LDH upon LPS/ATP and PolyI:C stimulation. This implies that the release of cytokines or inflammasomes by pyroptosis may be reduced upon Spermidine treatment. As recent data indicate that TLR4-mediated activation of microglia impairs microglial Aβ phagocytosis^44^, one may speculate that Spermidine ameliorates Aβ burden by preventing microglia from being constantly activated, i.e. switching to a DAM phenotype.

Most Spermidine actions are known to be attributed to its induction of autophagy^26,27^. Genetically mediated impairment of autophagy reversed the beneficial effects of Spermidine on longevity of worms, flies and yeast and its cardioprotective and anti-cancerous functions^26,29,37,60^. In accordance with these data, we show that Spermidine treatment of primary microglia and astrocytes increased the levels of the autophagic proteins BECN1 and LC3-II by reversing the inhibitory effect of LPS/ATP or PolyI:C on TFEB. TFEB is the main transcription factor for autophagy genes^61^ that also has been described as a Spermidine-regulated transcription factor in B cells^47^. Moreover, autophagy induction in glial cells by amino-acid starvation reduced cytokine levels similar to Spermidine and autophagy inhibition by 3-MA or BECN1 knockout blunted Spermidine-mediated reduction of cytokine release.

Previously, we used the opposite approach, showing that autophagy impairment by BECN1 reduction increased neuroinflammation and inflammasome activity in microglia^18^. Thus, our data underline our interpretation, namely that Spermidine-induced autophagy in glial cells accounts for the observed reduction of neuroinflammation. In support of this idea, several studies suggest an involvement of autophagic mechanisms in regulating AD. Fasting or caloric restriction is a strong activator of autophagy and was found to reduce Aβ deposition in different mouse models^4,62^. Rapamycin, an inhibitor of the mechanistic Target of Rapamycin (mTOR) and well-known activator of autophagy, not only prolonged the life span of several species similar to Spermidine but also ameliorated AD pathology in different AD mouse models, as reviewed by Kaeberlein et al.^63^. Likewise, treatment of AD-like mouse models with Metformin, another established inducer of autophagy, resulted in decreased Aβ pathology^64^, emphasizing the notion that the herein described effects of Spermidine in APPPS1 mice on Aβ burden are autophagy-mediated.

Based on the data presented here, we consider the body-endogenous substance Spermidine as a novel, most attractive therapeutic dietary supplement in AD as it substantially reduced synaptotoxic soluble Aβ and attenuated the AD-relevant neuroinflammation. Spermidine’s multifarious intracellular interference points in both primary microglia and astrocytes is of additional advantage, especially in light of its good tolerability and its uncomplicated way to be administered orally. Since Spermidine supplementation is used in humans, there is an obvious potential in testing Spermidine in AD patients. Supporting data were obtained in a first clinical trial in which older adults with subjective cognitive decline were treated with wheat germ extract rich in Spermidine. This treatment resulted in short-term improvement of memory performance^65,66^ and protected brain regions known to be affected in the course of AD^67^. Based on our data unraveling the mode of Spermidine’s action, the extension of Spermidine supplementation from individuals with subjective cognitive decline to clinical trials aimed at testing Spermidine efficacy in AD patients are thus justified.

## Materials and Methods

### Mice and Spermidine treatment

APPPS1^+/-^ mice^39^ were used as an Alzheimer’s disease-like mouse model. *Casp1^-/-^* mice were a kind gift from F. Knauf and M. Reichel, Medizinische Klinik m.S. Nephrologie und Internistische Intensivmedizin, Charité Berlin. Beclin1^flox/flox^ mice were a kind gift from Tony Wyss-Coray (Stanford University School of Medicine/USA).

APPPS1^+/-^ mice and littermate wild type control mice were treated with 3 mM Spermidine dissolved in their drinking water (changed twice a week) from an age of 30 days until an age of either 120 days or 290 days. Control mice received only water (H_2_O). Prior to each exchange of the drinking bottles, the weight of the bottles was determined and used to calculate the average volume consumed per animal per day. Animals were kept in individually ventilated cages with a 12 h light cycle with food and water *ad libitum*. All animal experiments were conducted in accordance with animal welfare acts and were approved by the regional office for health and social service in Berlin (LaGeSo).

### Tissue preparation

Mice were anesthetized with isoflurane, euthanized by CO_2_ exposure and transcardially perfused with PBS. Brains were removed from the skull and sagitally divided. The left hemisphere was fixed with 4 % paraformaldehyde for 24 h at 4°C and subsequently immersed in 30% sucrose until sectioning for immunohistochemistry was performed. The right hemisphere was snap-frozen in liquid nitrogen and stored at −80°C for a 3-step protein extraction using buffers with increasing stringency as described previously^68^. In brief, the hemisphere was homogenized in Tris-buffered saline (TBS) buffer (20 mM Tris, 137 mM NaCl, pH = 7.6) to extract soluble proteins, in Triton-X buffer (TBS buffer containing 1% Triton X-100) for membrane-bound proteins and in SDS buffer (2% SDS in ddH_2_O) for insoluble proteins. The protein fractions were extracted by ultracentrifugation at 100,000 g for 45 min after initial homogenization with a tissue homogenizer and a 1 ml syringe with G26 cannulas. The respective supernatants were collected and frozen at −80°C for downstream analysis. Protein concentration was determined using the Quantipro BCA Protein Assay Kit (Pierce) according to the manufacturer’s protocol with a Tecan Infinite^®^ 200 Pro (Tecan Life Sciences).

### Quantification of Aβ levels and pro-inflammatory cytokines

Aβ40 and Aβ42 levels of brain protein fractions were measured using the 96-well MultiSpot V-PLEX Aβ Peptide Panel 1 (6E10) Kit (Meso Scale Discovery, K15200E-1). While the TBS and TX fraction were not diluted, the SDS fraction was diluted 1:500 with PBS. Cytokine concentrations were measured in the undiluted TBS fraction using the V-PLEX Pro-inflammatory Panel 1 (Meso Scale Discovery, K15048D1). For all samples, duplicates were measured and concentrations normalized to BCA values.

### Histology

Paraformaldehyde-fixed and sucrose-treated hemispheres were frozen and cryosectioned coronally at 40 μm using a cryostat (Thermo Scientific HM 560) and stored afterwards in cryoprotectant (0.65 g NaH_2_PO_4_ × H_2_O, 2.8 g Na_2_HPO_4_ in 250 ml ddH_2_O, pH 7.4 with 150 ml ethylene glycol, 125 ml glycerine) at 4°C until staining. For immunohistochemistry, sections were washed with PBS and incubated with pentameric formyl thiophene acetic acid (pFTAA, 1:500, Sigma-Aldrich) for 30 min at RT. Subsequently, cell nuclei were counterstained with DAPI (1:2000, Roche, 10236276001) and sections embedded in fluorescent mounting medium (Dako, S3023). For quantification of pFTAA positive Aβ plaques, images of 10 serial coronal sections per animal were taken with an Olympus BX53 microscope, equipped with the QImaging camera COLOR 12 BIT and a stage controller MAC 6000 (1.25x objective). Images were analyzed using ImageJ by defining the cortex as the region of interest (ROI). Images were converted to grey scale and by using the same threshold for all sections, the pFTAA-positive area [in %] was obtained. Additionally, the average plaque size was determined and further analyzed by performing a plaque size distribution using thresholds for the size of the pFTAA-positive particles. The average of all 10 sections per animal was displayed in the graphs.

### Isolation and culture of adult microglia

Adult microglia were isolated from the hemispheres of 160 day old C57BL/6J mice by magnetic activated cell sorting (MACS). The manufacturer’s protocols were followed. In brief, mice were transcardially perfused with PBS and tissue dissociated with the Neural Tissue Dissociation kit (P) (Miltenyi Biotec, 130-092-628) in C-tubes (Miltenyi, 130-096-334) on a gentleMACS Octo Dissociator with Heaters (Miltenyi Biotec, 130-096-427). Afterwards, the cell suspension was labelled with CD11b microbeads (Miltenyi Biotec, 130-093-634) and passed through LS columns (Miltenyi Biotec, 130-042-401) placed on a OctoMACS™ manual separator. Subsequently, microglia were collected by column flushing and cultured in DMEM medium (Invitrogen, 41966-029) supplemented with 10% FCS (PAN-Biotech, P40-37500) and 1% penicillin/streptomycin (Sigma, P0781-20ML). Medium was changed every three days until adult microglia were treated as indicated after 8 days *in vitro* (DIV).

### Cell culture of neonatal microglia and astrocytes

Newborn mice (1-4 days old) were sacrificed by decapitation. Mixed glial cultures were prepared as described previously^18^. In brief, brains were dissected, meninges removed and brains mechanically and enzymatically homogenized with 0.005% trypsin/EDTA. Cells were cultured in complete medium consisting of DMEM medium (Invitrogen, 41966-029) supplemented with 10% FCS (PAN-Biotech, P40-37500) and 1% penicillin/streptomycin (Sigma, P0781-20ML) at 37°C with 5% CO_2_. From 7 DIV on, microglia proliferation was induced by adding 5 ng/ml GM-CSF (Miltenyi Biotec, 130-095-746) to the complete medium. Microglia were harvested at 10-13 DIV by manually shaking flasks for 6 min. Cells were treated after a settling time of 24 h. Neonatal BECN1^flox/flox^.CX3CR1^CreERT2^ microglia were treated with (Z)-4-Hydroxytamoxifen (Sigma #7904) and assessed 7 days after Tamoxifen treatment.

After isolating neonatal microglia, neonatal astrocytes were separated by MACS. Neonatal astrocytes were detached with 0.05% trypsin, pelleted by centrifugation and incubated with CD11b microbeads (Miltenyi Biotec, 130-093-634) for 15 min at 4°C to negatively isolate astrocytes. Afterwards, the cell suspension was passed through LS columns (Miltenyi Biotec, 130-042-401) placed on an OctoMACS™ manual separator and the flow-through containing the astrocytes was collected. Subsequently, astrocytes were cultured in complete medium for 2 days before being treated. For all experiments, 100,000 cells were seeded on 24 well plates if not stated otherwise.

### Cell treatment

For pro-inflammatory stimulation, cells were either treated with LPS (1 μg/ml, Sigma, L4391-1MG) for 3 h followed by ATP (2 mM, Sigma Aldrich, A6419-5g; 4 mM for ASC speck/inflammasome formation) for 45 min or with PolyI:C (50 μg/ml, InVivoGen, tlrl-picw-250) for 6 h if not stated otherwise. Spermidine trihydrochloride (Sigma, S2501-5G) diluted in complete medium was added as indicated. Autophagy was activated by keeping cells in HBSS for 2 h prior to treatment (24020-091, Invitrogen) and blocked by addition of 3-MA (Sigma-Aldrich, M9282, final concentration 10 mM). The ASC oligomerization inhibitor MCC950 (inh-mcc, Invivogen) was used with a final concentration of 300 nM.

### Western blot

For ASC crosslinking, 1 mM DSS (Thermo, A39267) was added to freshly harvested microglia in PBS for 30 min. All cell pellets were lysed in protein sample buffer containing 0.12 M Tris-HCl (pH 6.8), 4% SDS, 20% glycerol, 5% β-mercaptoethanol. Proteins were separated by Tris-Tricine polyacrylamide gel electrophoresis (PAGE) according to Schägger and Jagow 1987^69^ and transferred by wet blotting onto a nitrocellulose membrane. After blocking with 1% skim milk in Tris-buffered saline with 0.5% Tween20 (TBST), the following primary antibodies were used: BECN1 (1:500, Cell Signaling, 3495), LC3 (1:500, Sigma, L8918), CASP1 and pro-CASP1 (1:500, Abcam, ab179515), IL-1β and pro-IL-1β (1:500, eBioscience, 88701388), NLRP3 (1:500, AdipoGen, AG-20B-0014), Gasdermin D (1:500, Adipogene, AG-25B-0036-C100), ASC (1:500, Adipogene, AG-25B-0006-C100) and ACTIN (1:10,000, Sigma, A1978). For signal detection the SuperSignal West Femto Maximum Sensitivity Substrate (ThermoFisher, 34096) was used. Western blots were analyzed by quantifying the respective intensities of each band using ImageJ. All samples were normalized to ACTIN levels or whole protein content in the supernatant.

### Quantitative real-time PCR

For total RNA isolation, the RNeasy Mini kit (Qiagen, 74104) was used and cells were directly lysed in the provided RLT lysis buffer in the cell culture plate. Reverse transcription into cDNA was performed using the High-Capacity cDNA Reverse Transcription kit (ThermoFisher, 4368813). The manufacturer’s instructions for both kits were followed. Quantitative PCR was conducted on a QuantStudio 6 Flex Real-Time PCR System (Applied Biosystems) using 12 ng cDNA per reaction. Gene expression was analysed in 384 well plates by using the TaqMan Fast Universal Master Mix (Applied Biosystems, 4364103) and TaqMan primers for *β-Actin* (ThermoFisher, Mm00607939_s1), *Il-6* (ThermoFisher, Mm00446190_m1), *Tnf-α* (ThermoFisher, Mm00443258_m1), *Nlrp3* (ThermoFisher, Mm00840904_m1), *Casp1* (ThermoFisher, Mm00438023_m1), *Tfeb* (ThermoFisher, Mm00448968_m1). Within the Double delta Ct method, values were normalized to the house keeping gene *Actin* and nontreated controls.

### ELISA

Cytokine concentrations in the supernatant of cultured cells were measured using the IL-1β (eBioscience, 88701388), TNF-α (eBioscience, 88723477) and IL-6 (eBioscience, 88706488) enzyme-linked immunosorbent assay (ELISA) kit according to manufacturer’s instructions. The absorption was read at a wavelength of 450 nm and a reference length of 570 nm with the microplate reader Infinite^®^ 200 Pro (Tecan Life Sciences) and analyzed using the Magellan Tecan Software.

### Cytotoxicity assay

Cell cytotoxicity was detected by measuring lactate dehydrogenase (LDH) levels using the CytoTox 96^®^ Non-Radioactive Cytotoxicity Assay (Promega, G1780). The manufacturer’s instructions were followed and the absorbance reflecting the LDH content in the cell supernatant was measured at 492 nm (600 nm reference) with an Infinite^®^ 200 Pro (Tecan Life Sciences) plate reader. All values were normalized to non-treated controls.

### Immunocytochemistry and Confocal Microscopy

Cells were seeded at a density of 50,000 cells per well on 12 mm coverslips. After treatment, cells were fixed with 4% paraformaldehyde for 20 min, permeabilized with 0.1% Triton X-100 in PBS for another 20 min and blocked with 3% bovine serum in PBS for 1 h. The primary antibody anti-ASC (AdipoGen, AG-25B-0006, 1:500) was added overnight at 4°C. Subsequently, cells were incubated with the fluorescent secondary antibodies Alexa Fluor 568-conjugated anti-rabbit IgG (1:500, Invitrogen, A11011) for 3 h at RT. Cell nuclei were counterstained with DAPI (Roche, 10236276001) and coverslips embedded in fluorescent mounting medium (Dako, S3023). Images were acquired using Leica TCS SP5 confocal laser scanning microscope controlled by LAS AF scan software (Leica Microsystems, Wetzlar, Germany). Z-stacks were taken and images presented as the maximum projection of the z-stack. The number and size of ASC specks was assessed using ImageJ software as described before^18^.

### Data analysis

All values are presented as mean ± SEM (standard error of the mean). For pairwise comparison between two experimental groups, the student’s t-test was used. Statistical differences between more than two groups were assesses with One-way ANOVA using the indicated *post hoc* test. Statistically significant values were determined using the GraphPad Prism software and are indicated as follows: *P < 0.05, **P < 0.01 and ***P < 0.001.

## Supporting information

All supplementary data

## Acknowledgement

This work was supported by the Deutsche Forschungsgemeinschaft (DFG, German Research Foundation) under Germany’s Excellence Strategy—EXC-2049—390688087, as well as under JE-278/6-1 to MJ and SFB TRR 43, SFB TRR 167 and HE 3130/6-1 to F.L.H., by the German Center for Neurodegenerative Diseases (DZNE) Berlin, and by the European Union (PHAGO, 115976; Innovative Medicines Initiative-2; FP7-PEOPLE-2012-ITN: NeuroKine). We are grateful to Alexander Haake for excellent technical support. Treatment images were created with Biorender.com.

## Author contributions

KF, SS, FH and MJ designed experiments; KF, NS and JH treated mice and analyzed the brains; KF, NS and LF performed experiments and analyzed data for the TLR3 pathway and astrocytes; JS, JH and MJ performed experiments and analyzed data for the TLR4 pathway; KF, NS, JS and MJ assessed autophagy; KF prepared the figures. All authors wrote, revised and approved the manuscript.

## Notes

### Competing Interest Statement

The authors have declared no competing interest.

