## Supplementary material for "The autophagy activator Spermidine ameliorates Alzheimer’s disease pathology and neuroinflammation in mice": All supplementary data

#### Supplementary Figure 1

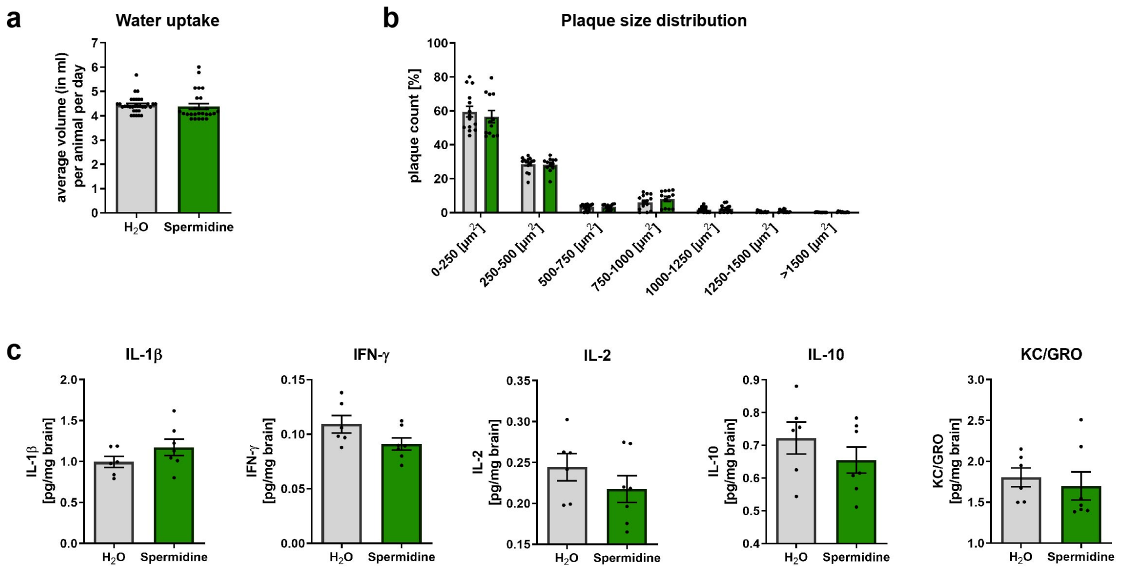


**Supplementary Figure 1**

APPPS1 mice were treated with 3 mM Spermidine via their drinking water and analysed at an age of 290 days. Spermidine-treated APPPS1 mice were compared to non-treated controls ($H_2O$).

(**a**) The average volume in ml per animal per day was calculated based on the volume that was drank in an interval of 3.5 days per cage. APPPS1 $H_2O$ (n = 32), APPPS1 Spermidine (n = 26).

(**b**) Tissue sections were stained for Aβ plaques with pFTAA and a plaque size distribution of the cortex was calculated. The number of plaques of the indicated sizes is displayed as a percentage of the total number of plaques detected. APPPS1 $H_2O$ (n = 14), APPPS1 Spermidine (n = 12).

(**c**) The content of the cytokines IL-1β, IFN-γ, IL-2, IL-10 and KC/GRO in the TBS fraction of brain homogenates of male Spermidine-treated APPPS1 mice was measured using electrochemiluminescence (MesoScale Discovery panel). APPPS1 $H_2O$ (n = 6), APPPS1 Spermidine (n = 7).

Mean ± SEM, two-tailed t-test, * $p < 0.05$, ** $p < 0.01$.

#### Supplementary Figure 2

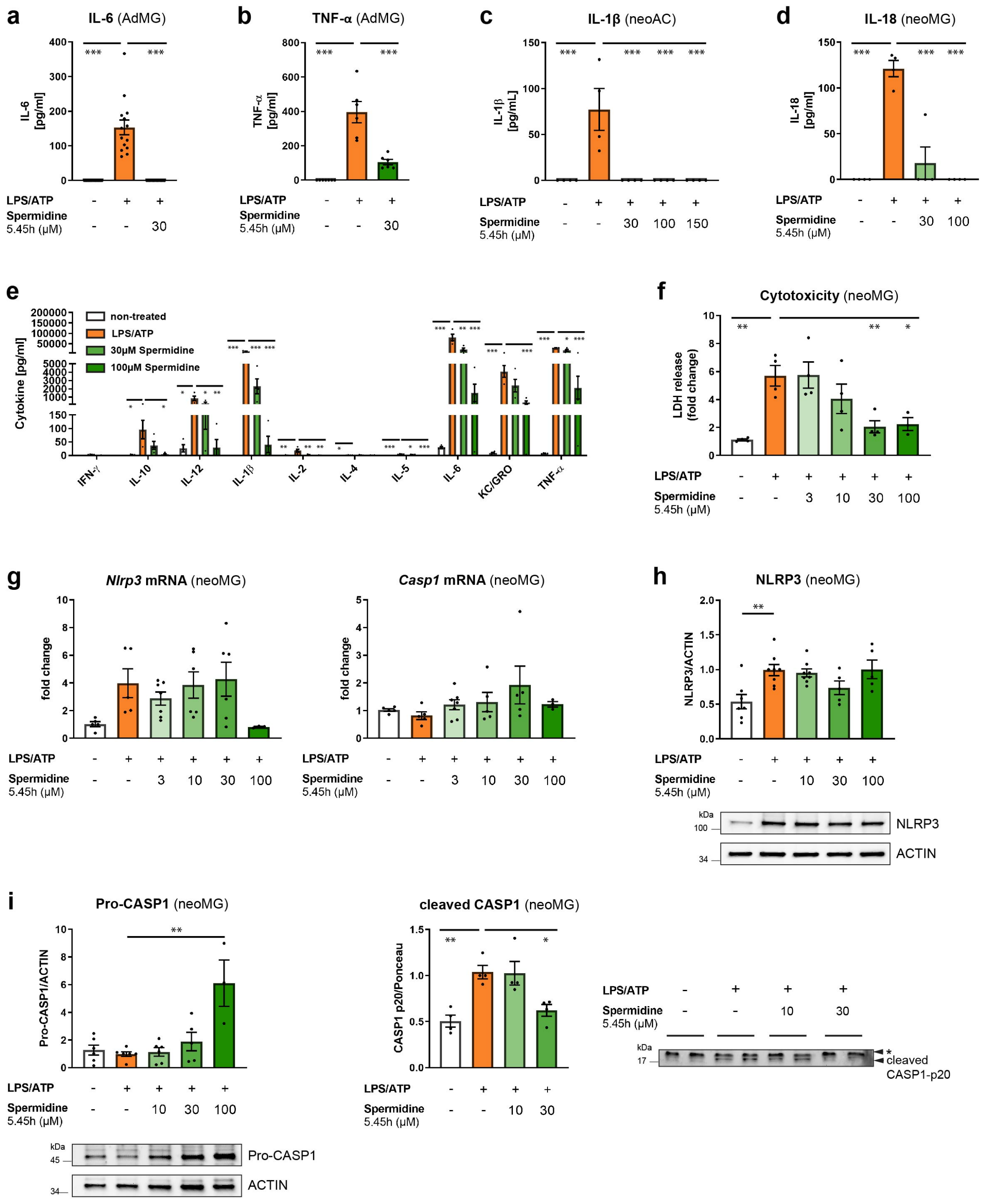

#### Supplementary Figure 2

Neonatal microglia (neoMG), adult microglia (AdMG) and neonatal astrocytes (neoAC) were treated with LPS/ATP and Spermidine as shown in Fig. 2a.

(**a**) The IL-6 concentration in the cell supernatant of adult microglia was determined by ELISA; n = 14-15.

(**b**) The TNF-α concentration in the cell supernatant of adult microglia was determined by ELISA; n = 6-7.

(**c**) The IL-1β concentration in the cell supernatant of neonatal astrocytes was determined by ELISA; n = 4.

(**d**) The IL-18 concentration in the cell supernatant of neonatal microglia was determined by ELISA; n = 4.

(**e**) The amount of cytokines in the cell supernatant was determined by electrochemiluminescence (MesoScale Discovery panel); n = 3-7.

(**f**) The cytotoxicity of Spermidine was determined by measuring LDH release into the cell supernatant. LDH release was normalized to the non-treated control; n = 3-4.

(**g**) The gene expression of *NLRP3* and *Pro-Casp1* was assessed by RT-qPCR. Their expression was normalized to *Actin* and displayed as fold change compared to non-treated control cells; n = 3-7.

(**h**) NLRP3 protein levels were determined by Western blot and normalized to ACTIN. Representative images are shown and values are displayed as fold changes compared to LPS/ATP-treated cells; n = 5-8.

(**i**) Pro-CASP1 protein levels and cleaved CASP1 p20 protein levels were determined by Western blot and normalized to ACTIN or whole cell protein (Ponceau C). Representative images are shown (* non-specific band) and values are displayed as fold changes compared to LPS/ATP-treated cells; n = 3-6.

Mean ± SEM, one-way ANOVA, Dunnett´s post hoc test (reference= LPS/ATP-treated cells), * $p < 0.05$, ** $p < 0.01$, *** $p < 0.001$.

#### Supplementary Figure 3

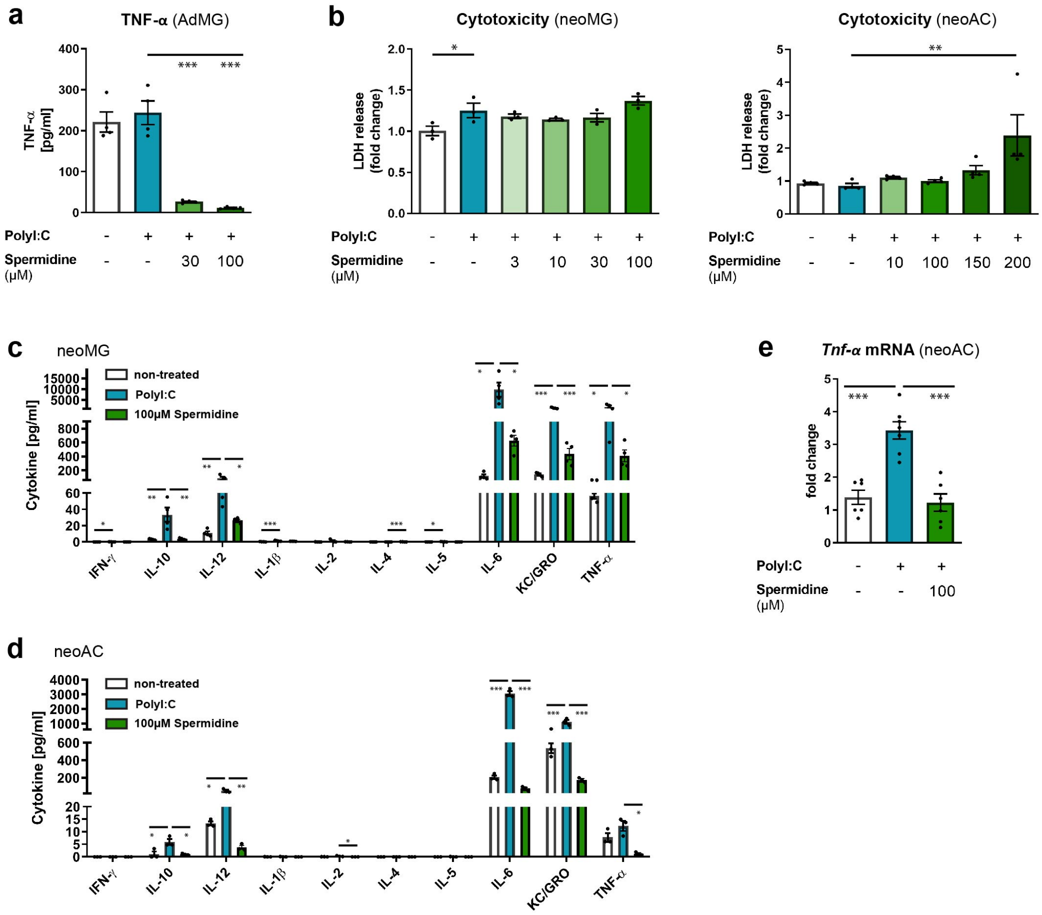


#### Supplementary Figure 3

Neonatal microglia (neoMG), adult microglia (AdMG) and astrocytes (neoAC) were stimulated with PolyI:C (50 µg/ml) and the indicated concentrations of Spermidine for 6 h according to the treatment scheme in Fig. 3a.

(**a**) The concentration of TNF-α in the cell supernatant of adult microglia was measured by ELISA; n = 4.

(**b**) The cytotoxicity of Spermidine was determined by measuring LDH release into the cell supernatant of neonatal microglia and astrocytes. LDH release was normalized to the non-treated control; microglia: n = 3; astrocytes: n = 4.

(**c-d**) The concentrations of the cytokines IL-6, TNF-α, IL-10, IL-12 and KC/GRO in the cell supernatant of stimulated (**c**) neonatal microglia and (**d**) astrocytes were determined using electrochemiluminescence (MesoScale Discovery panel); microglia: n = 3; astrocytes: n = 3.

(**e**) The gene expression of *Tnf-α* was assessed by RT-qPCR. Its expression was normalized to *Actin* and displayed as fold change compared to non-treated control astrocytes. n = 3.

Mean ± SEM, one-way ANOVA, Dunnett´s post hoc test (control = PolyI:C-treated cells), * $p < 0.05$, ** $p < 0.01$, *** $p < 0.001$.

### Supplementary Figure 4

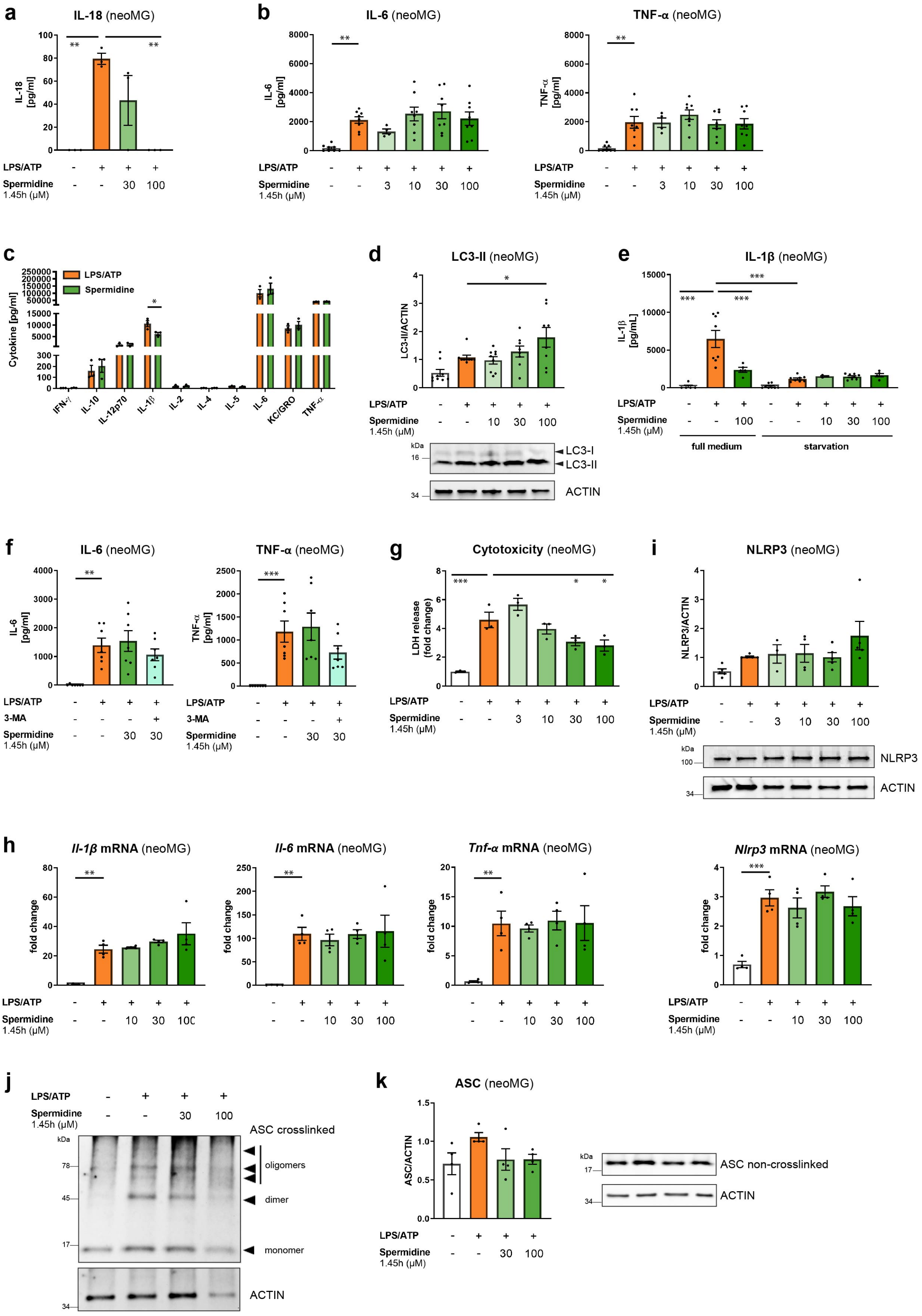

**Supplementary Figure 4**

Neonatal microglia (neoMG) were treated with LPS/ATP and Spermidine as shown in Fig. 5a.

**(a)** The IL-18 concentration in the cell supernatant was determined by ELISA; n = 3.

**(b)** The IL-6 and TNF-α concentration in the cell supernatant was determined by ELISA; n = 4-8.

**(c)** The amount of cytokines in the cell supernatant was determined by electrochemiluminescence (MesoScale Discovery panel); n = 3.

**(d)** LC3-II levels were determined by Western blot and normalized to ACTIN. Representative images are shown and values are displayed as fold changes compared to LPS/ATP controls; n = 8-9.

**(e)** Neonatal microglia were stimulated as in Fig. 5a either in full medium (the same data points as in Fig. 4e) or in starvation medium HBSS (non-treated and LPS/ATP are the same data points as in Fig. 4e) and the IL-1β concentration in the cell supernatant was determined by ELISA; n = 3-8, one-way ANOVA, Tukey´s post hoc test.

**(f)** Neonatal microglia were stimulated as in Fig. 5a and 3-MA was added simultaneously with Spermidine. The IL-6 and TNF-α concentration in the cell supernatant was determined by ELISA; n = 7.

**(g)** The cytotoxicity of Spermidine was determined by measuring LDH release into the cell supernatant of neonatal microglia. LDH release was normalized to the non-treated control; n = 3.

**(h)** The gene expression of *Il-1β, Tnf-α* and *Il-6* was assessed by RT-qPCR. The expression was normalized to *Actin* and displayed as fold change compared to non-treated control cells; n = 4.

**(i)** The gene expression of *Nlrp3* was assessed by RT-qPCR. Its expression was normalized to *Actin* and displayed as fold change compared to non-treated control cells; n = 4. NLRP3 protein expression was determined by Western blot and normalized to ACTIN. Representative images are shown and values are displayed as fold changes compared to LPS/ATP-treated cells; n = 3-5.

**(j)** Proteins were chemically crosslinked by DSS. ASC monomer and oligomers were visualized by Western blot.

(**k**) ASC monomer protein expression was determined by Western blot and normalized to ACTIN. Representative images are shown and values are displayed as fold changes compared to LPS/ATP-treated cells; $n = 4$.

Mean ± SEM, one-way ANOVA, Dunnett´s post hoc test (reference = LPS/ATP-treated cells) if not stated otherwise, * $p < 0.05$, ** $p < 0.01$, *** $p < 0.001$.
